# peaks2utr: a robust Python tool for the annotation of 3’ UTRs

**DOI:** 10.1101/2022.05.26.493605

**Authors:** William Haese-Hill, Kathryn Crouch, Thomas D. Otto

**Affiliations:** Institute of Infection, Immunity & Inflammation, MVLS, University of Glasgow, Glasgow, UK

## Abstract

**Summary:** Annotation of non-model organisms is an open problem, especially the detection of untranslated regions (UTRs). Correct annotation of UTRs is crucial in transcriptomic analysis to accurately capture the expression of each gene yet is mostly overlooked in annotation pipelines. Here we present peaks2utr, an easy-to-use Python command line tool that uses the UTR enrichment of single-cell technologies, such as 10x Chromium, to accurately annotate 3’ UTRs for a given canonical annotation.

**Availability and Implementation:** peaks2utr is implemented in Python 3 (≥ 3.8). It is available via PyPI at https://pypi.org/project/peaks2utr and GitHub at https://github.com/haessar/peaks2utr. It is licensed under GNU GPLv3.

## Introduction

Despite advances in genome assembly, little progress has been made in annotation, notwithstanding the vital role annotations play in the functional interpretation of data. The popular annotation pipeline MAKER2 is estimated to predict ~80% of genes correctly (Holt and Yandell, 2011), highlighting the challenge in sequence annotation. Moreover, the annotation of UTR regions is generally ignored in most non-model organisms (Huang and Teeling, 2017). But the 3’ UTRs of messenger RNAs (mRNAs) are known to regulate mRNA-based processes, such as mRNA localization, mRNA stability, and translation (Mayr, 2019). Thus, understanding the expression and differential usage of 3’ UTRs is important in functional inference. But comprehensive genome annotation pipelines neglect to offer annotation of 3’ UTRs, for example, this functionality was not implemented in Companion (Steinbiss, et al., 2016) because UTR annotations in reference genomes were generally not good enough for model training at the time of publication. But to fully understand a species and how genes are regulated, fully annotated genomes are required. Further, for comprehensive RNA-seq analysis, reads from the UTR should be captured for analysis. This is especially true for methods where most reads map toward the end of UTRs, like 10x Chromium (Wang, et al., 2021).

To improve this situation, we have developed peaks2utr to update 3’ UTR models in existing genome annotations using data from 10x Chromium sequencing runs, where signal is inherently concentrated at the distal ends of the 3’ or 5’ UTRs, allowing concise inference of UTR boundaries. The method described here addresses the use of 3’ 10x Chromium sequencing to improve 3’ UTR annotation. Its method is precise, as it seeks to apply a soft-clipped polyA-tail truncation (SPAT) algorithm to read pileups. We will show that peaks2utr is more accurate, as well as easier to install and use, than existing tools (Huang and Teeling 2017) and demonstrate the utility of its application in different datasets.

## Workflow

peaks2utr (implemented in Python) is called from the command line and takes as input a GFF/GTF annotation file (with or without existing three prime UTR annotations) and a BAM file containing aligned read data, as well as various optional parameters (see Supplementary Text).

BAM files are split into forward and reverse strands, and further into read groups, using pysam (a Python wrapper for htslib and SAMtools (Danecek, et al., 2021)). Now the SPAT algorithm is applied: reads containing soft-clipped bases and polyA-tails of a given length are detected, and their end bases tallied as “truncation points” (see Supplementary Text).

Peaks are called from the BAM file using MACS2 (Zhang, et al., 2008). Each peak is iterated over, and a search is performed for genes falling within a user-defined maximum distance of base pairs from the peak. Subsequently, a peak is designated a 3’ UTR if it passes a set of criteria and truncated using SPAT if possible (Figure 1, Supplementary Text).

**Figure 1:**
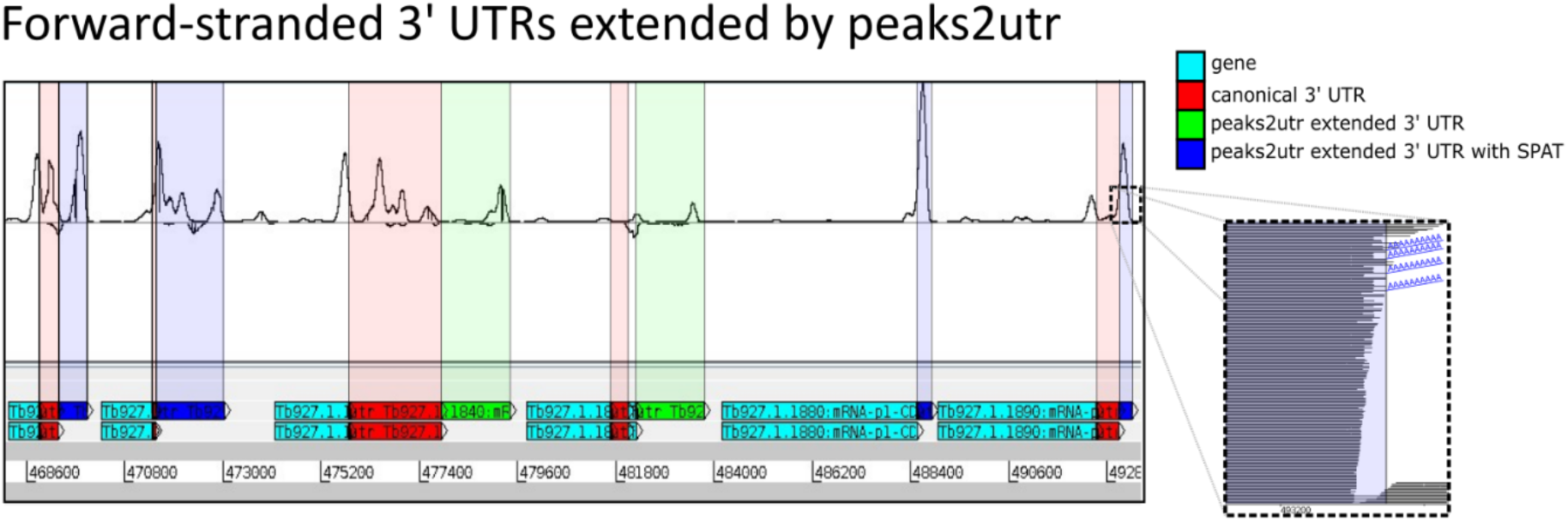
*Trypanosoma brucei* TREU927 genes (teal) with canonical 3’ UTR (red). Top track shows the same genes with extension by peaks2utr (green and blue), and how this coincides precisely with coverage peaks in 3’ region; only the fourth gene from the left saw no extension – peaks2utr matched within a few bases. Inset shows magnification of mapped read stack, where SPAT has been applied to multiple reads with shared end base.

Although MACS2 (version 2.2.7.1), is optimised for ChIP-seq data, it was found to sufficiently predict regions of peaks likely to correspond to a UTR when tuned for “broad peaks”.

## Application

To test the performance of peaks2utr, we applied it to *Caenorhabditis elegans* reference chromosome I, which has manually curated 3’ UTRs. We compared our results to GETUTR (Kim, et al., 2015) a recent, well-cited tool for UTR annotation that takes the same inputs as peaks2utr. Overall, peaks2utr was able to find more UTRs matching those in the canonical annotation (see Supplementary Table 1). Further, peaks2utr did not overpredict the UTR length, which is in part due to the implementation of SPAT. A comparison of the two sets of new UTRs called by both tools revealed that 85 of them matched, with a number of these worthy candidates for future annotation (see Supplementary Figure 2).

Next, we applied peaks2utr to the parasite *Trypanosoma brucei* (Briggs, et al., 2021). Overall, out of 11,703 annotated genes, 6,179 have 3’ UTR annotations. By applying peaks2utr we obtain in total 9,907 3’ UTRs, an increase of ~60% and covering ~85% of available genes. Of these, 3,728 are new, 5,349 are altered and the remaining 830 are unchanged. Figure 1 shows an example of 7 annotated genes with 3’ UTRs extended by peaks2utr. For the 12.4 GB BAM file of ~140 million mapped reads, and using 12 processors, runtime was approximately 50 minutes; GETUTR was unable to run for a BAM file this large.

To understand the impact of the changed annotation against the canonical reference annotation, we re-ran Cell Ranger 6.1.1 with the updated reference from peaks2utr. The peaks2utr annotation saw an improvement in genes captured per cell of 889 from 632 (~33% increase), and UMI counts per cell of 1,111 from 797 (~39% increase), implying that a substantial signal is gained by using improved 3’ UTR annotations from peaks2utr.

## Discussion

Here we presented peaks2utr that generates 3’ UTR annotations. It is easy to install, maintainable and includes unit testing. Leveraging the capture of the polyA-tail of the transcript allows us to map the transcript end to an unprecedented precision. Interestingly, existing tools do not perform as well (Supplementary Table 1) and struggle to scale with the sheer amount of data (see Supplementary Text). We demonstrated that although peaks2utr is not perfect, it does a substantially better job than comparable tools. With the advent of scRNA-seq and its wide usage of non-model organisms, we recommend that scientists should run peaks2utr for their single cell experiments to improve the capture of the signal (see Figure 1). Further, it will improve the annotation of many species so far lacking 3’ UTR regions.

## Funding

Wellcome Trust funded WHH and TDO (104111/Z/14/Z & A) and KC 218288/Z/19/Z.

## Notes

### Competing Interest Statement

The authors have declared no competing interest.

