## Supplementary Methods for "peaks2utr: a robust Python tool for the annotation of 3’ UTRs"

Peaks2UTR is implemented in Python 3 (≥ 3.8). It is available via PyPI at https://pypi.org/project/peaks2utr and GitHub at https://github.com/haessar/peaks2utr. It is licensed under GNU GPLv3. Here we describe some of the implementation and test performed.

### Peak to UTR assignment criteria

For each MACS2 peak, a set of five criteria are applied to determine whether it corresponds to a valid UTR. Supplementary Figure 1 shows visual examples of each of these criteria being applied.

i) If peak falls within a user-defined max-distance of base-pairs from the 3’-end of any gene, then it will be considered a potential UTR.

ii) If peak intersects a 3’ UTR that is already annotated in the input GFF file, either ignore, override or extend, depending on user preference.

iii) If peak range is a subset of a gene, then ignore.

iv) If peak intersects the 5’-end of the following gene on the same strand, any potential UTR will be truncated to the start base of that gene.

v) If peak is within max-distance (see (i)) of a gene, but also corresponds to 3’-end of the following gene, any potential UTR will only be assigned to the following gene.

*Supplementary Figure 1 – Examples of peak assignment criteria; pink shaded regions correspond to MACS2 broad peak, green features are (potential) UTRs, teal features are genes.*

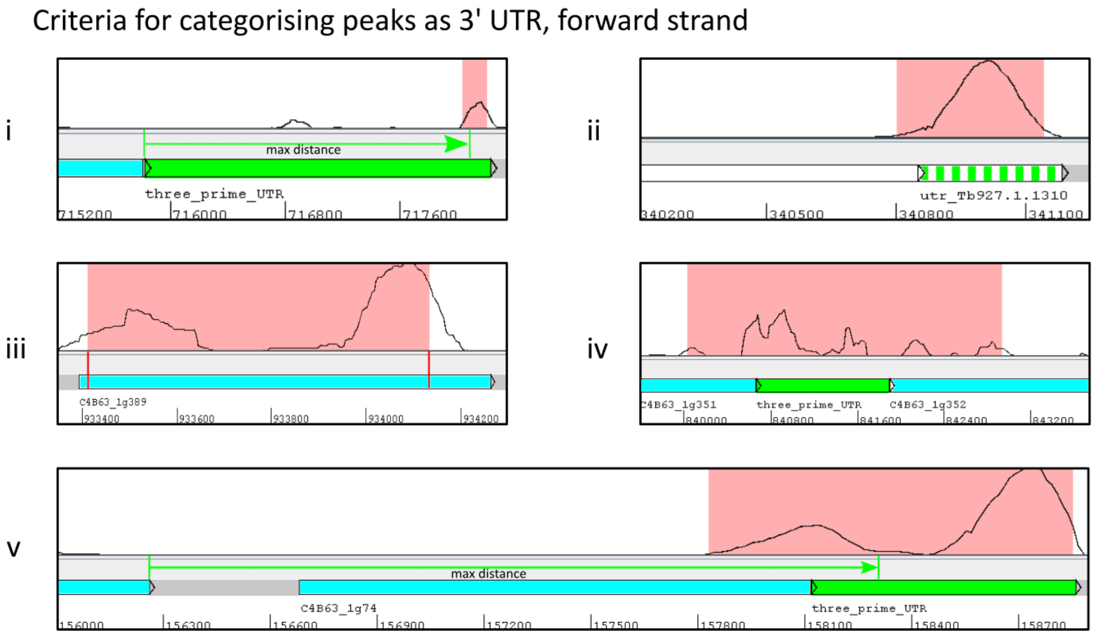

### Soft-clipped polyA-tail truncation (SPAT) algorithm

During pre-processing of BAM files, each read is parsed to determine whether it contains soft-clipped bases, and if so, whether the soft-clipped segment contains a polyA/T-tail with a minimum length specified by the user in an optional command line (CL) option --min-poly-tail (default 10). The mapped "end" bases of qualifying reads are tallied, and any that fall below a threshold specified by the user in an optional CL option --min-pileups (default 10) are filtered out.

The resulting set of bases are referred to as "truncation points": If any potential UTR passes the previously defined criteria, and a truncation point falls within its range, the UTR will be truncated to that base (or the outermost base in the case of multiple truncation points).

### Runtime enhancements

A substantial factor in the overall runtime of peaks2utr is the pre-processing of files:

- gffutils builds a sqlite3 database to contain feature models for the input GFF file.
- pysam splits input BAM file into multiple batches for each strand to guarantee strand-independence of peaks, as the previously defined criteria are applied differently for forward and reverse. Each of these resulting BAM files are indexed.
- SPAT algorithm outputs truncation points to json files.
- MACS2 calls peaks on stranded-BAM files and outputs to .broadPeak files.

To improve runtimes, peaks2utr harnesses multi-processing and caching of these pre-processed files (so that subsequent runs with tweaked parameters are significantly faster). The user can specify available processor cores with optional -p or –processors command line (CL) options. peaks2utr can utilise the inherent multi-processing functionality of samtools when calling any pysam function to manipulate BAM files, while gffutils and MACS2 pre-processing functions, being independent, are handled asynchronously.

SPAT pre-processing uses one available processor core for each BAM file split by strand/read group. MACS2 peaks for UTR annotation are batched depending on available processor cores.

The user must specify that the cache should be persisted with the –keep-cache CL option, to prevent any unwanted storage overhead.

As well as the three_prime_UTR annotations, a summary statistics text file is produced, informing the user about total UTR count, as well as number of peaks failing each criterion.

### *C. Elegans* application / GETUTR comparison

GETUTR offers 3 smoothing algorithms to remove "noise" in the RNA-seq signal: Max.fit and Min.fit preference local maxima and minima, respectively, while PAVA uses weighted least-squares regression. Each UTR is output as a fragmented series of features with an assigned normalization scaling *s* between 0 and 1.

We applied both GETUTR (version 2.0.0) and peaks2utr to *Caenorhabditis elegans* reference chromosome I , obtained from WormBase release WS283 (Davis, et al., 2022). 10x reads from a recent study (Packer, et al., 2019), with accession code GSE126954, were mapped against the full reference using 10x Genomics Cell Ranger 6.1.1 to generate the input BAM file. Reads were acquired from 12 runs of SRA experiment SRX5411289. The resulting BAM file was 19 GB in size, which reduced to 3.3 GB when filtering for only chromosome I mapped reads, as used for the test.

GETUTR was run with PAVA smoothing and all features with $s < 0.95$ were removed to ensure highest quality and minimise overprediction of UTRs. To prevent GETUTR from crashing due to excessive virtual memory usage, we had to adapt the source code to prevent reading into memory of the entire samfile and instead use an iterator.

peaks2utr was run with parameters --max-distance 2500, --override-utr, --min-pileups 4, and --min-poly-tail 5. Results of the comparison are summarised in Supplementary Table 1 and Supplementary Figure 2.

*Supplementary Table 1* – Comparison of peaks2utr with GETUTR on *C. Elegans*. Reported are different cases of missed, new or extended UTR by both tools compared to the canonical annotation. peaks2utr outperforms GETUTR, especially by not overpredicting the length.

** by more than 50 bp*
*** by more than 200 bp*

| | peaks2utr | | GETUTR | |
|---|---|---|---|---|
| | num | % | num | % |
| Missed UTR | 434 | 16.74 | 568 | 21.91 |
| Novel UTR | 143 | 6.21 | 151 | 6.94 |
| **UTR matches (within 50 bp)** | **773** | **33.58** | **164** | **7.54** |
| Shortened UTR* | 48 | 2.09 | 485 | 22.29 |
| Extended UTR* | 992 | 43.09 | 138 | 6.34 |
| **Too long UTR**** | **346** | **15.03** | **1238** | **56.89** |

*Supplementary Figure 2 – C. Elegans protein coding gene WBGene00020556 (teal) with novel 3' UTR as predicted by both peaks2utr (blue) and GETUTR (red), one of 85 novel UTRs predicted by both tools.*

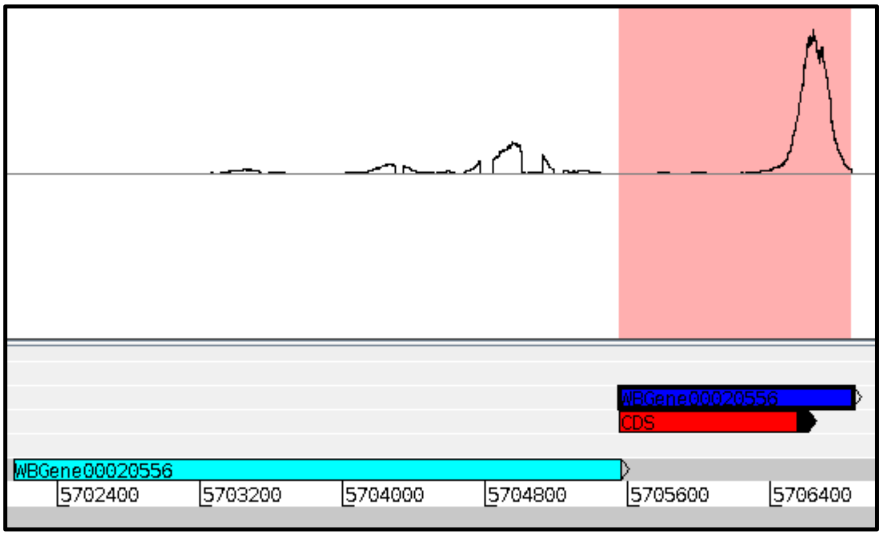


### *T. Brucei* application

For both runs, we ran 10x Genomics Cell Ranger 6.1.1 using 10x reads that were obtained from European Nucleotide Archive with study accession number PRJEB41744 and experiment accession number ERX5428965. The canonical annotation *Trypanosoma brucei brucei* TREU927 reference was obtained from TriTrypDB, release 43 (Aurrecoechea, et al., 2016).

peaks2utr was run with parameters --max-distance 2500, --override-utr.
